## Supplementary Figures for "Spectral cues are necessary to encode azimuthal auditory space in the mouse superior colliculus"

Supplementary Figure 1, Ito et al., 2019

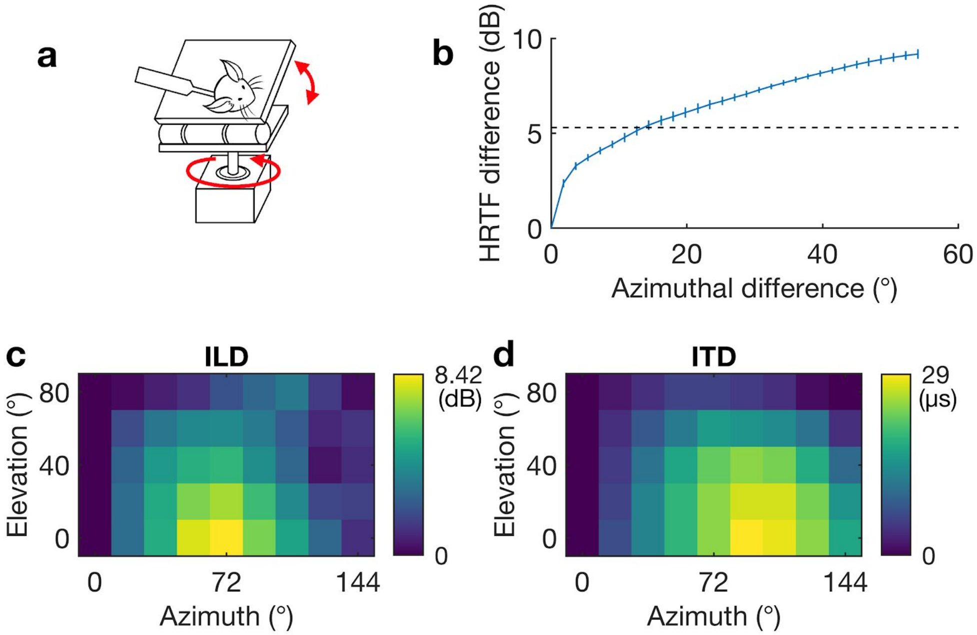


Supplementary Figure 1: Head-related transfer function (HRTF) measurement and virtual auditory stimulation.

a: A schematic of the stage for HRTF measurement. A mouse head with a microphone attached to the back of the ear canal is fixated on the stage. The stage has automated control of rotation and manual control of tilt, allowing the measurement in the entire quadrant of the auditory sensory field without uncoupling the microphone from the ear canal. Foam materials are padded on all of the hard surfaces of the stage to prevent sound reflection.

b: Average difference of the HRTFs as a function of the difference of the incident azimuth. The average difference of HRTFs was evaluated by taking the root mean square (RMS) errors of two HRTFs with a fixed difference of their incident azimuthal angles and averaging them over the frequency and the incident angle. The HRTF differences increase monotonically as a function of the difference of the incident azimuth. The black dashed line indicates the RMS difference of the HRTFs between different animals (3 pairs from 3 animals). We estimated the angular error due to the use of non-individual HRTF is ~±14°.

c: ILDs at 45 points for the right-top quadrant of the mouse. The values for the ipsilateral side are a mirror image of these values with an opposite sign. The ILD at each point is the difference in the average intensity of the stimulus between 5 kHz and 80 kHz. A cubic interpolation of ILDs in the horizontal plane identifies ~65° as the azimuth that maximize the ILD.

d: ITDs at 45 points for the right-top quadrant of the mouse. ITDs were measured by the time difference between the peaks of the head-related impulse response (time domain expression of the HRTF) of left and right ears.

Supplementary Figure 2, Ito et al., 2019

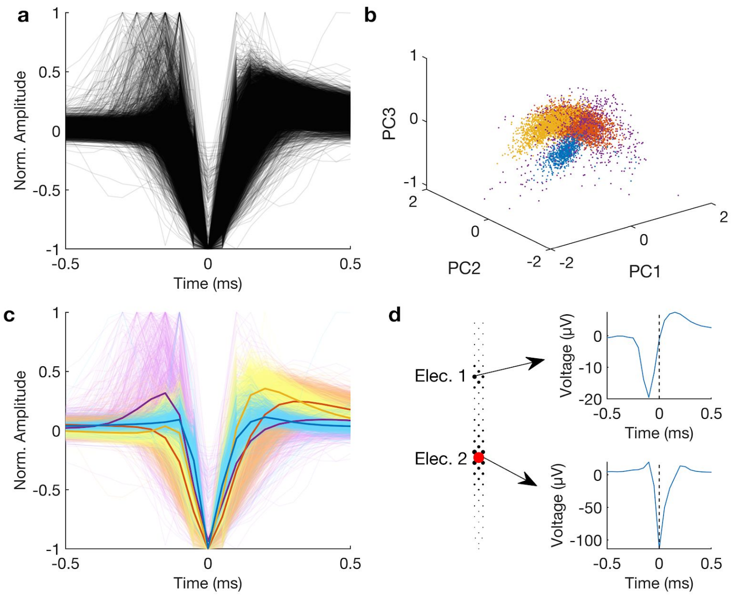


Supplementary Figure 2: Separation of the axonal signals.

a: Overlaid average spike waveforms for all 6060 neurons in the dataset.

b: A 3 dimensional scatter plot of the waveforms in the space of the 3 most important principal components. The color of the dots indicates the result of the clustering analysis based on a mixture of Gaussians model. Note that the purple cluster is larger and sparser than the others (the determinant of the Gaussian covariance matrix (i.e. volume) of the four clusters were $7.6 \times 10^{-5}$, $7.1 \times 10^{-5}$, $1.7 \times 10^{-6}$, and $1.3 \times 10^{-2}$, for the yellow, red, blue, and purple clusters, respectively, showing that the purple cluster is more than two orders of magnitude larger than the second-largest cluster). It indicates that the purple cluster is fitting to outliers. These outliers contain mostly biphasic spikes with opposite polarity (a peak followed by a trough) that are detected at the dendrites near the soma. Therefore we did not exclude this cluster from the analysis.

c: Same as (a) with each waveform colored with the corresponding cluster color in (b). The bold solid lines indicate the average waveform of each cluster. Of these 4 traces, the blue color indicates the axonal signal, based on the shape of the waveform that is consistent with an axonal signal (see (d)). These neurons were excluded from the analysis.

d: Example waveforms of the somatic signals and axonal signals detected from one neuron. The circles on the left panel indicate the amplitudes of the spike of this neuron detected by electrodes at the corresponding locations. The red circle indicates the electrode where this neuron was identified in spike-sorting. The average voltage traces of this neuron at two electrodes were compared on the right panels. The spike at Electrode 1 appears 0.1 ms earlier than that at Electrode 2, indicating that the spike on Electrode 2 is an axonal signal of the neuron whose cell body is located near Electrode 1. With such observations, we concluded that a spike with a right shoulder (yellow and orange trances in (c)) is a cell body signal, and a narrow and triphasic signal with a weaker right shoulder (blue traces in (c)) is an axonal signal.

Supplementary Figure 3, Ito et al., 2019

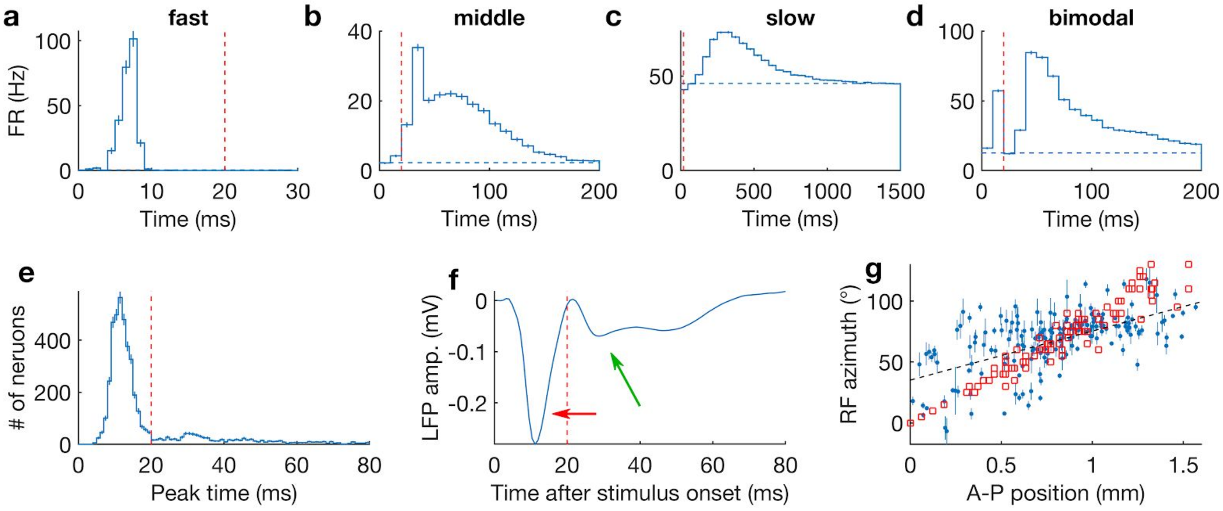


Supplementary Figure 3: Temporal responses of the auditory responsive neurons.
a, b, c, d: Example temporal responses of four neurons with different time scales (fast, middle, slow, and bimodal). The red dashed line indicates 20 ms, the division line for fast and slow responses. Note that the time scale of each figure is largely different. The error bars are estimated by Poisson statistics.
e: Histogram of the peak response time (the time of the bin with the highest firing rate) for all neurons that had significant responses within 100 ms from the stimulus onset. The peak response time is bimodally distributed, suggesting that the activity before and after ~20 ms can be separated.
f: Average stimulus-evoked local field potentials (LFPs) that show bimodal temporal responses. These LFPs were measured on an electrode with the maximum evoked LFP amplitude. The average LFP time course was calculated based on individual LFPs of all the virtual locations and trials. After the stimulus onset, two troughs were observed (red and green arrows). These timings are, again, separated at 20 ms and coincide with the peak responses of individual neurons (e). Because the LFPs are largely generated by synaptic inputs and the return current associated with it[40], this result suggests that the SC receives temporally segregated inputs before and after 20 ms from the stimulus onset.
g: The map of auditory space constructed by the responses of all neurons with a localized receptive field (RF) in the 20–100 ms time scale. The blue dots show the auditory RF azimuths versus the physical locations, along the A–P axis, of the individual neurons when solely looking at the spikes in the 20–100 ms range; the red squares are the visual RF azimuths measured by multiunit activity in the superficial SC. The slope and offset of the auditory RFs are 40 ± 5 °/mm and 35 ± 4°, respectively. The correlation of the physical A–P positions and RF azimuth is weaker ($r = 0.59$) than that for the fast time scale ($r = 0.70$).

Supplementary Figure 4, Ito et al., 2019

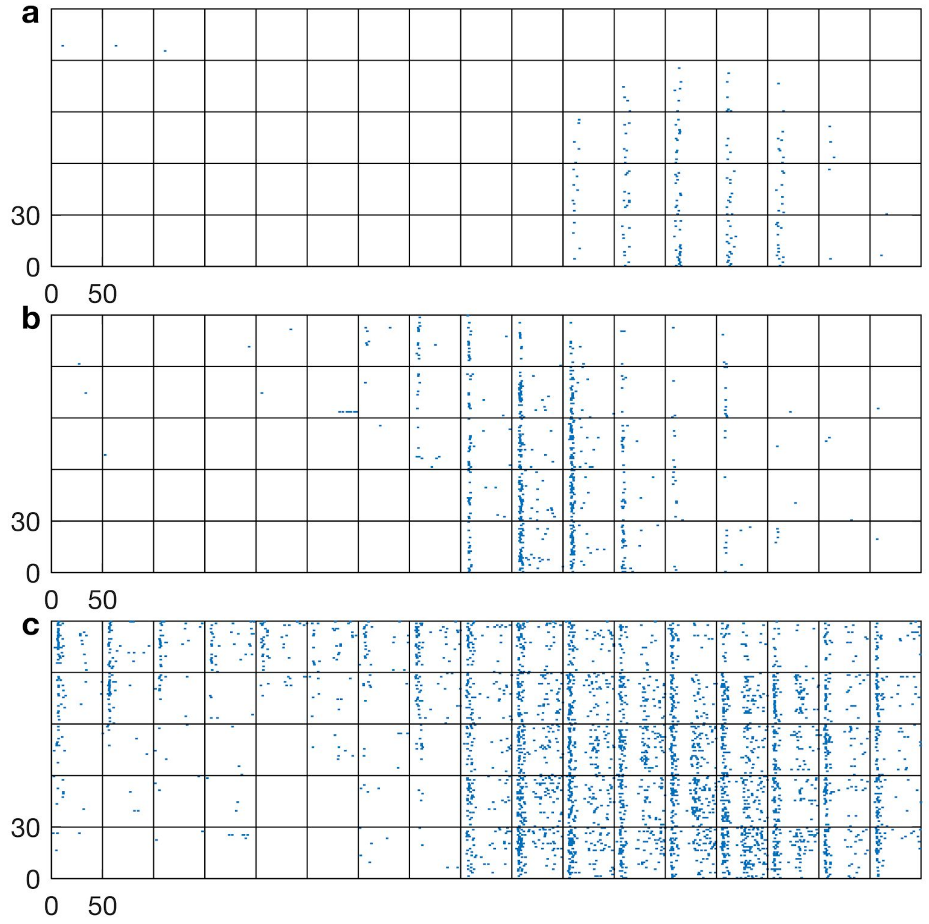


Supplementary Figure 4: Raster plots of the neurons that are shown in Fig. 2a-c. The individual panels in the figure correspond to each of the 85 virtual sound source directions. In each panel, the spike timings (ms) are plotted for 30 stimulus repetitions.

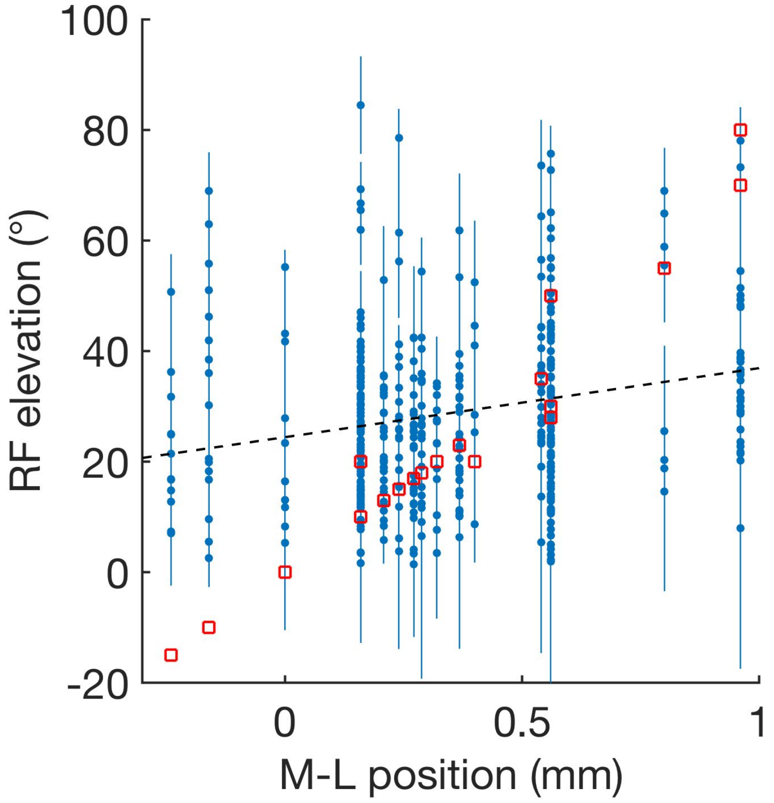


Supplementary Figure 5: A scatter plot of mediolateral (M–L) SC positions vs. RF elevations showing topographic organization along the M–L axis of the SC. The exploratory dataset and the combined dataset had significant slopes (18 ± 6 °/mm and 13 ± 5 °/mm, respectively), but the blinded dataset alone did not have a significant slope (5 ± 7 °/mm; see Blind analysis subsection in Methods). The plotted dataset is the combined dataset. Red squares are visual RFs measured by multiunit activity in the superficial SC. The black dashed line is the $\chi^2$ fit to the data including systematic errors (see Methods for details). The slope of the auditory map is much smaller than that of the visual map along the elevation axis (75 ± 4 °/mm; measured by multi-unit activity in the superficial SC). The correlation coefficient of the combined data is r = 0.22.

Supplementary Figure 6, Ito et al., 2019

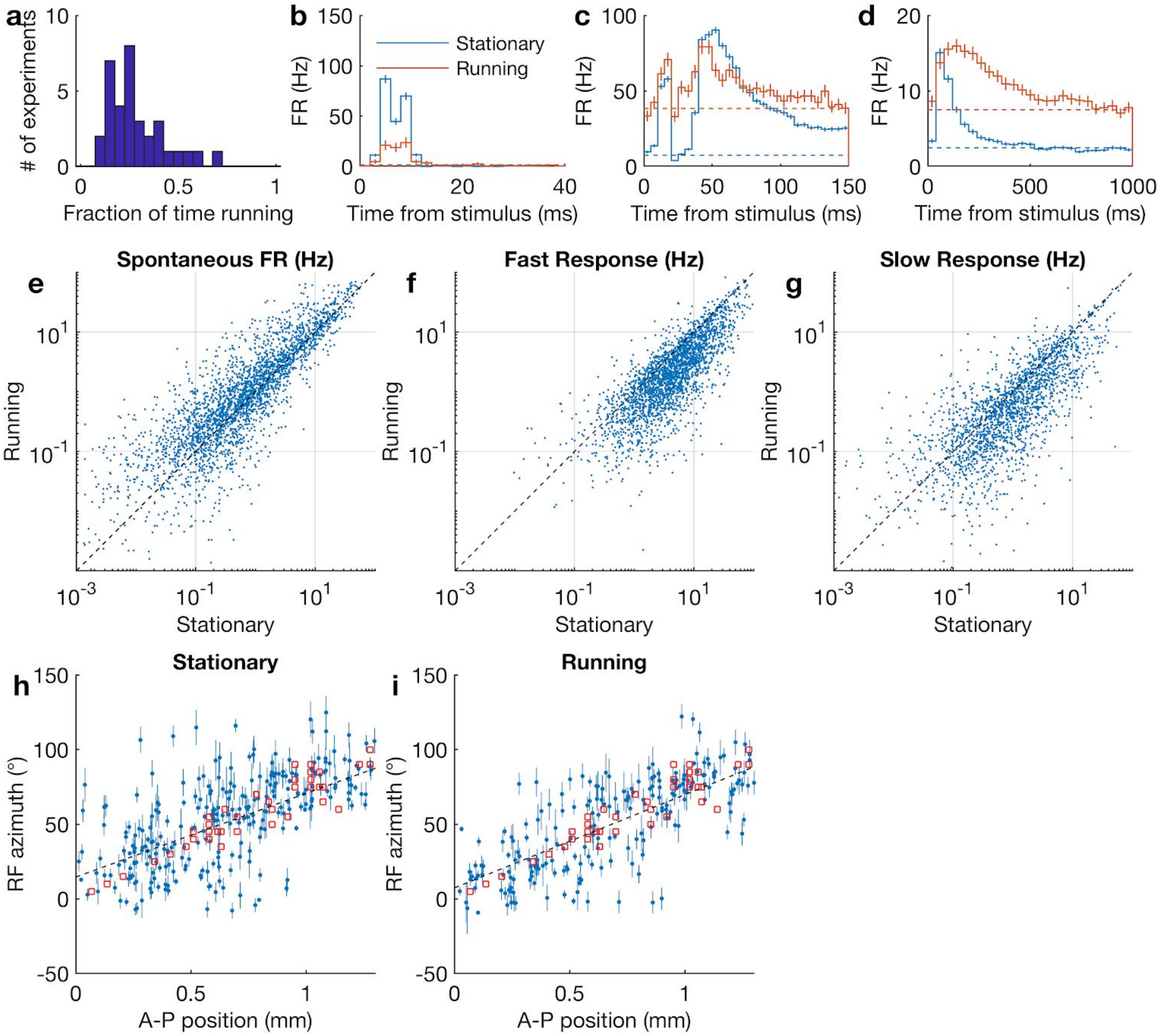


Supplementary Figure 6: Effect of locomotion on the auditory responses of SC neruons.
a: Histogram of the fraction of running time (running speed >1 cm/s) in each of the 31 experiments.
b–d: Temporal responses of the neurons during stationary periods (running speed ≤1 cm/s) and running periods at different time scales. The dashed lines indicate the spontaneous FRs; the full lines indicate the sum of the auditory response and spontaneous FRs. Enhancement of the spontaneous firing rate during running is observed in (c) and (d); suppression of the evoked firing rate (the histogram above the dashed line) during running is observed in (b) and (c). A change in the temporal response pattern during running was observed in (d).
e–g: Summary scatter plot of the firing rate changes caused by locomotion. Spontaneous firing rate increases and both fast and slow auditory responses are suppressed by locomotion.
h, i: The auditory map of space based on spikes during the stationary (h) or running (i) periods showing no change in the parameters of the topographic map dependent on locomotion. The slope and offset for the stationary topographic map are 56 ± 4 °/mm and 14 ± 3°; those for the running topographic map are 62 ± 4 °/mm and 8 ± 4°.

Supplementary Figure 7, Ito et al., 2019

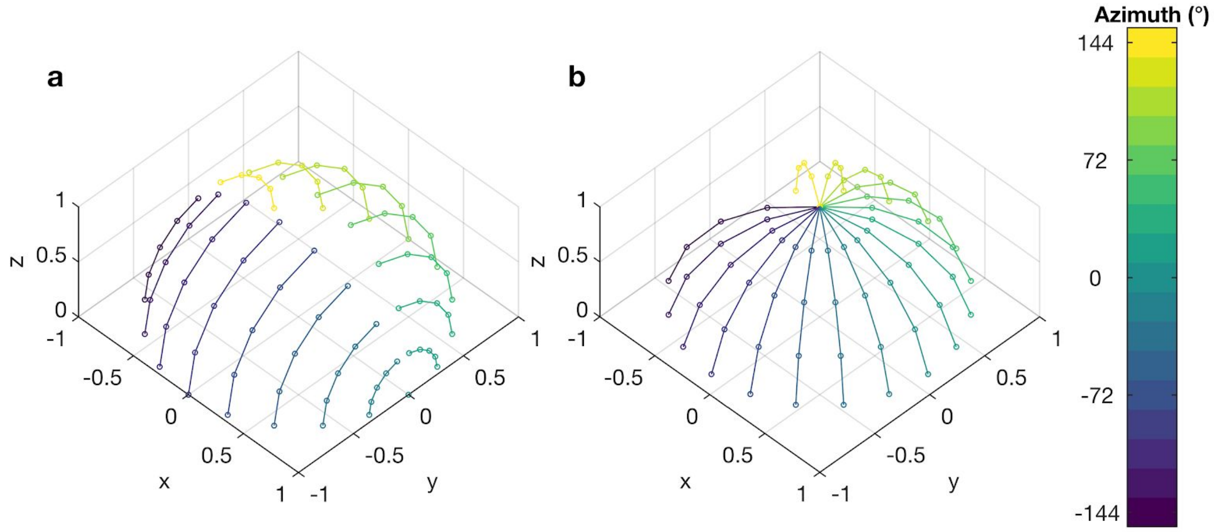


Supplementary Figure 7: Two definitions of the spherical polar coordinate system.
a: Polar coordinate system with its Z-axis defined as the anterior direction of the animal (Front-Z coordinate). This coordinate system was used to measure the HRTFs, to present the stimuli to the animal during experiments, and to plot the RF heat maps that were used in Figs. 1d–f, 2a–j, 3a–j, 4a, and 4d–g.
b: Polar coordinate system with its Z-axis defined as the dorsal direction of the animal (Top-Z coordinate). This coordinate system was used to fit a Kent distribution to the experimental data and extract the RF azimuth from the neuronal responses. By using this coordinate system, we avoided the issue of having a spurious azimuthal value associated with the pole at the front.

Supplementary Figure 8, Ito et al., 2019

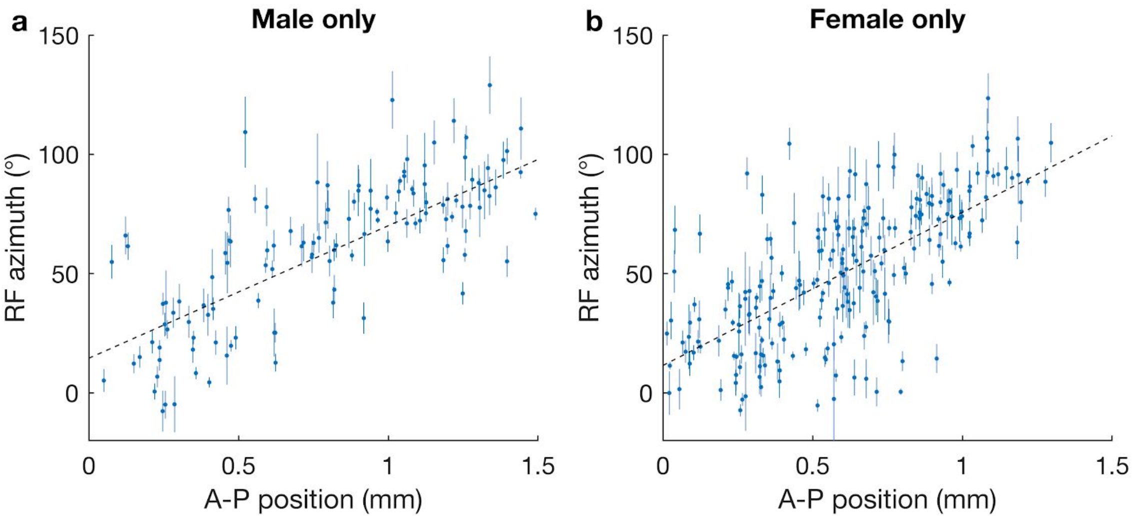


Supplementary Figure 8: Consistent map properties between male and females.
a: Same topographic map structure shown in Fig. 2e, but used only male data. The slope and offset are 55 ± 5 °/mm and 15 ± 5 °.
b: Similar to a, but used only female mice. The slope and offset are 64 ± 5 °/mm and 12 ± 4 °. These are not significantly different from those of males (p = 0.23 and 0.59, respectively).
